# A neural network model delivers a highly prognostic protein signature in cancer stem cells that identifies relapse in stage III colorectal cancer patients

**DOI:** 10.64898/2026.01.12.697945

**Authors:** Anna Sturrock, Sanghee Cho, Manuela Salvucci, Marc Sturrock, Joanna Fay, Tony O’Grady, Elizabeth McDonough, Christine Surrette, Jinru Shia, Canan Firat, Nil Urganci, Batuhan Kisakol, Emer P. O’Connell, John P. Burke, Niamh M. McCawley, Deborah A. McNamara, John F. Graf, Simon S. McDade, Mohammadreza Azimi, Daniel B. Longley, Fiona Ginty, Jochen H. M. Prehn

## Abstract

**Background:** Stage III colorectal cancer poses a significant threat of metastasis development, as tumour resection and adjuvant chemotherapy do not guarantee prolonged disease-free survival.

**Objective:** The spatial, quantitative, and qualitative characteristics of various cell types within tumour tissues could be key to developing accurate prognostic AI models.

**Design:** Tissue microarrays created from primary tumour tissues collected during surgical resection from a cohort of 493 stage III colorectal cancer (CRC) patients were analysed for 61 protein markers at the single-cell level using multiplexed immunofluorescence imaging via the Cell DIVE™ platform. Subsequent cell-type classification enabled quantitative cell-type analyses, co-localisation neighbourhood assessments, and cell-type-specific protein signature discoveries that distinguish between early and late/non-recurring patient samples.

**Results:** This study identifies a stem cell protein profile that drives tumour relapse. A deep neural network (DNN) model, based on a stem cell protein signature composed of BAX, MLKL, FLIP, GLUT1, and CDX2, provided accurate prognosis for stage III CRC patients in both discovery and validation cohorts and in an independent validation cohort. Nodal count-based metric further increased prognosis accuracy. Our study also revealed distinct spatial arrangements of immune, endothelial, and stem cells that were linked to early tumour recurrence.

**Conclusion:** Our findings propose a clinically promising prognostic tool based on a five-protein stem cell signature. These markers not only predict chemotherapy resistance in cancer stem cells but also suggest potential therapeutic strategies such as combinatorial treatments incorporating small molecule inhibitors targeting FLIP and GLUT1.

**Key messages:** *What is already known on this topic:*

- More than 20% of stage III colorectal cancer patients will experience early tumour recurrence within the first 3 years post treatment that includes surgery and adjuvant 5-FU based chemotherapy treatment.
- Several studies pointed towards involvements of number of cell type specific spatial neighbourhoods in tumour progression where some immune tumour microenvironment promoting angiogenesis and intravasation events, some may provide immunosuppression.
- Cancer stem cells could be responsible for metastatic tumour spread, early recurrence and chemoresistance.

*What this study adds:*

- Spatial single cell quantitative multiplex profiling of 45 cancer hallmark proteins and 15 cell identity markers in 493 stage III CRC patients tissue samples demonstrated significant differences in cellular proximity neighbourhoods, cell type specific abundance and expression between the early and late recurrence samples.
- We discover that macrophages show spatial association with the blood vessels in early recurrence samples. Moreover, we observed conglomeration of B cells and macrophages with Tregulatory, Thelper and Tcytotoxic cells in association with early recurrences.
- We showed that stromal abundance of Tregulatory, Thelper, Tcytotoxic cells and monocytes are significantly in late, and no recurrence samples compared to early recurrence samples.
- The most differential expression profile that differentiates late and no recurrence samples from the early recurrence samples is related to the stem cell population. Particularly, we found overexpression of GLUT1, FLIP and downregulation of BAX, BAK, MLKL and CDX2 proteins in the cancer stem cell of early recurrence samples.
- We built a neural network based on the cancer stem cell protein signature (BAX, MLKL, FLIP, GLUT1 and CDX2 proteins) that delivers a high-performance prognostic classifier.

*How this study might affect research, practice or policy:*

- Our results propose a clinically promising prognostic tool based on a five-protein stem cell signature that outperforms existing clinical and proposed transcriptomic based signatures for separation between risk groups.
- Moreover, our five-protein signature markers not only predict stem cell chemotherapy resistance and therefore tumour recurrence but also suggest potential therapeutic strategies. For instance, this approach could guide combinatorial treatments at high risk of chemoresistance, such as incorporating small molecule inhibitors targeting FLIP (currently in discovery phase) and GLUT1 (already under preclinical trial evaluation).

## Introduction

Colorectal cancer (CRC) is the second leading cause of cancer-related death worldwide. The demographics of this disease have changed dramatically over the last decade, with surging numbers of patients being detected before the age of 40, resulting in an estimated 73% increase in CRC related deaths by 2040 according to the World Health Organization (2023, July 11).

The advanced stages of CRC are defined by the spread of cancer cells into lymph nodes and potentially other parts of the body. Stage III CRC is characterized by the spread of micrometastatic cells into lymph nodes surrounding the primary tumour site. This spread is believed to be initiated by a population of pluripotent cancer epithelial cells including cancer stem cells (CSCs) that are the most efficient “seeders” of metastasis due to their high clonogenic capacity ^1^. In addition to metastasis, CSCs also contribute to tumour growth and chemotherapy resistance ^2^. The current standard-of-care for stage III treatment involves surgical resection of the primary tumour and positive lymph nodes, followed by adjuvant oxaliplatin/fluoropyrimidine/leucovorin (5-fluorouracil (5FU), FOLFOX; or xeloda/capecitabine, XELOX) chemotherapy. However, around 24% of stage III patients will experience tumour recurrence within the first 3 years post-surgery ^3^. It is believed that many of these patients will progress because of the persistence of disseminated CSCs enabled by their resistance to adjuvant chemotherapy.

In this study, we investigated the spatial, quantitative and qualitative characteristics of CSCs and other cell types in treatment-naive stage III tumours resected from the patients followed for at least 3 years post-surgery. Tissue microarrays (TMAs) were constructed from tumour tissues collected during surgical resection of primary tumours in a cohort of 493 stage III CRC patients who underwent adjuvant chemotherapy (Figure 1). We quantified proteomic markers with multiplexed immunofluorescence imaging performed with the Cell DIVE™ platform followed by image processing, single cell quantification and cell type classification. We evaluated at the single cell level 61 protein markers, including markers for cell segmentation and identification, epithelia/stroma/endothelial segregation, cell proliferation and differentiation, microsatellite stability, epithelial mesenchymal transition (EMT), apoptosis and necroptosis, metabolism and immune signalling. We performed comparisons between frequencies of cell types, the expression of cell ‘state’ markers in individual cell types, and their spatial proximity and explored how these parameters correlate with early recurrence. Analysing separate discovery, validation and independent validation cohorts, we identify the existence of a highly prognostic CSCs specific protein signature. Based on this new signature, we developed a deep neural network (DNN) model that facilitates robust prognosis of recurrence.

**Figure 1.**
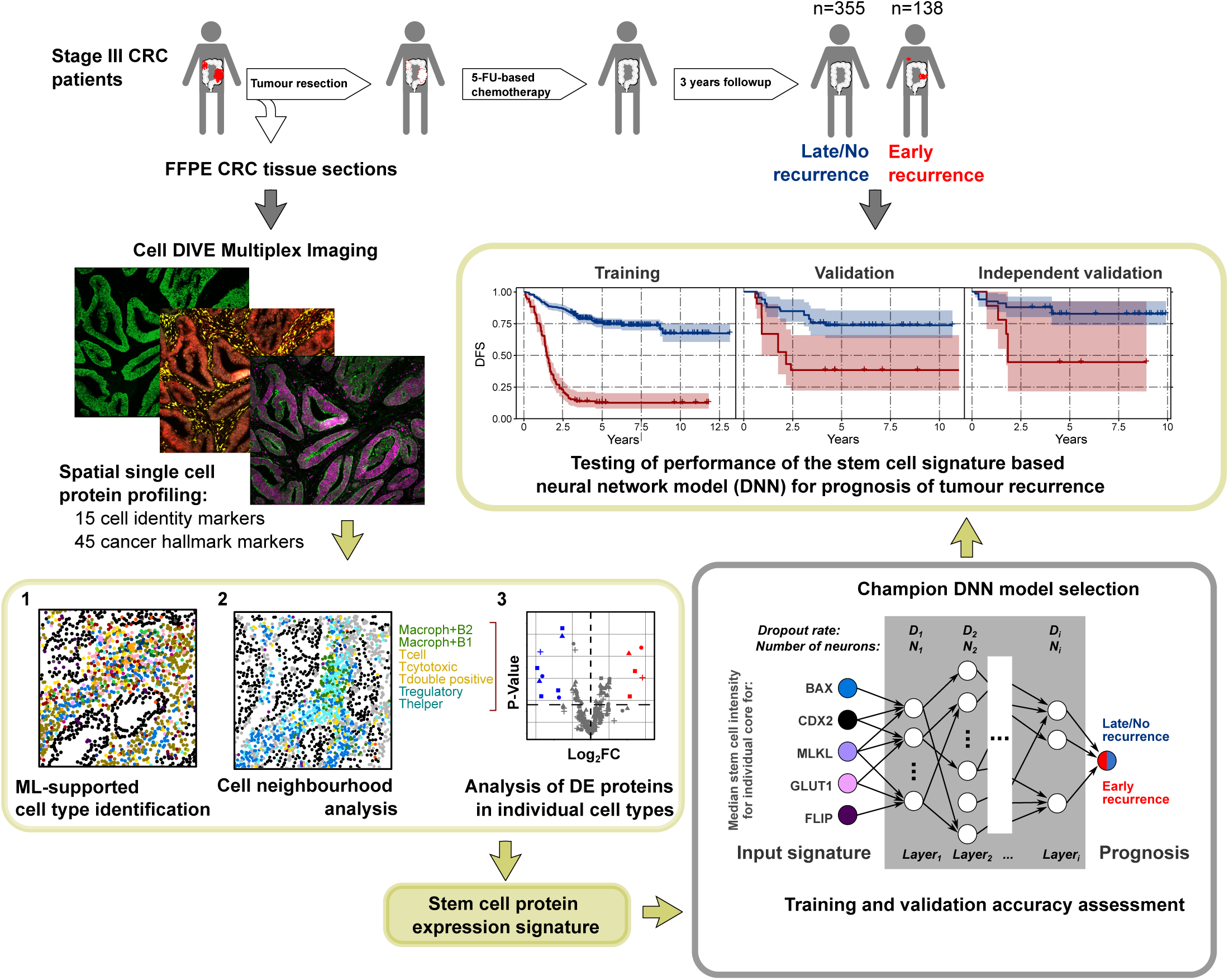
Schematic diagram representing the design of this study.

## Results

### Robust cell type classification pipeline

We developed a pipeline for cell type identification for high-plex single cell data analysis in stage III tumour tissue (*see Methods*). The pipeline begins with the recognition of epithelial CDX2^+^, PCK26^+^ and AE1^+^ positive cells (‘cancer cells’). Secondly, the pipeline identifies Monocytes (CD3^−^CD8^−^CD4^+^), T regulatory cells (CD3^+^CD4^+^FOXP3^+^), T helper cells (CD3^+^CD4^+^FOXP3^−^), double negative T cells (T cell CD3^+^CD8^−^CD4^−^FOXP3^−^), cytotoxic T cells (CD3^+^CD8^+^) and double positive T cells (CD3^+^CD4^+^CD8^+^) (**Figure 2A**). Thirdly, macrophages (Macrophage, CD4↑CD68↑), B cell-associated macrophages (Macroph+B1, CD4↑CD68↑CD20↑, Macroph+B2, CD4↑CD68↑modCD20) and endothelial cells (Endothelial_1, CD31↑CD34↑COLIV↑SMA↑, Endothelial _2, CD31↑CD34↓COLIV↑SMA↑) are classified (**see Supplementary Methods**). As the final step, the pipeline classifies the stem cell (ALDH1↑KI67↓AE1↓) population (**Figure 2B**).

**Figure 2.**
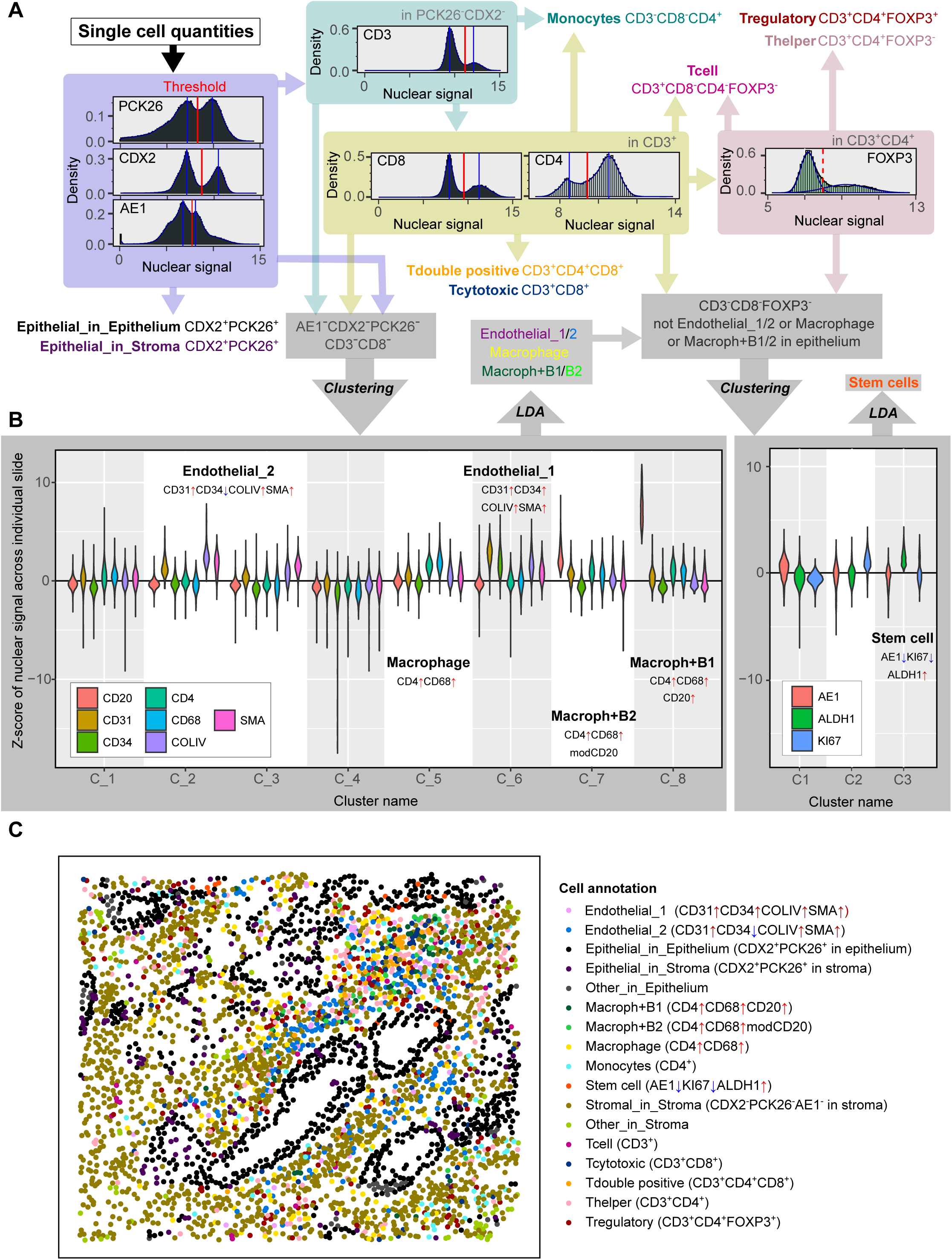
Illustration of cell type classification pipeline. (A) Automated single binary marker threshold detection algorithms determined epithelial and T cells over TMA slides. (B) Algorithm for multiple marker dependent detection of endothelial, macrophage and stem cells rely on hierarchical clustering following linear discriminant analysis (LDA) based classification model. (C) The example of final cell annotation illustrated on one of the cores.

Final cell type annotations (**Figure 2C**) established on the individual TMA slides ensured an accurate, unbiased and unsupervised classification, compared to, for example, a manually supervised annotation affected by a quantitatively limited training set, or deep learning-based classification pipelines that could be affected by model overfitting ^4^ or experimental batch effects.

### Stem, immune and endothelial cells proximity neighbourhoods differentiate early from late/non-recurring patients

Recent advances in tissue spatial proteome multiplex technologies (for example, Cell DIVE™, GeoMx® or CODEX ^5^ in conjunction with a range of spatial analysis computational tools ^6–10^ have accelerated the discovery of cellular neighbourhoods that dictate the course of CRC progression. Therefore, we first investigated the spatial cellular landscape and potential differences between early and late/non-recurring tumour samples from all four clinical cohorts (**Table S1**). We calculated the co-localization neighbourhood probabilities and correlated them between different cell types using the hoodscanR package ^8^. These correlation coefficients were collated into the median coefficients over two groups of cores: early and late/non-recurrence. The final pairwise median coefficients were clustered to recognise the cell type-specific neighbourhoods that are potentially associated with propensity of recurrence (**Figure 3A, B**). We tested proximity neighbourhoods of different sizes and chose the optimal proximity of 20 nearest cells in the final co-localisation probabilities calculation that showed the most discriminant features between the two groups of cores.

**Figure 3.**
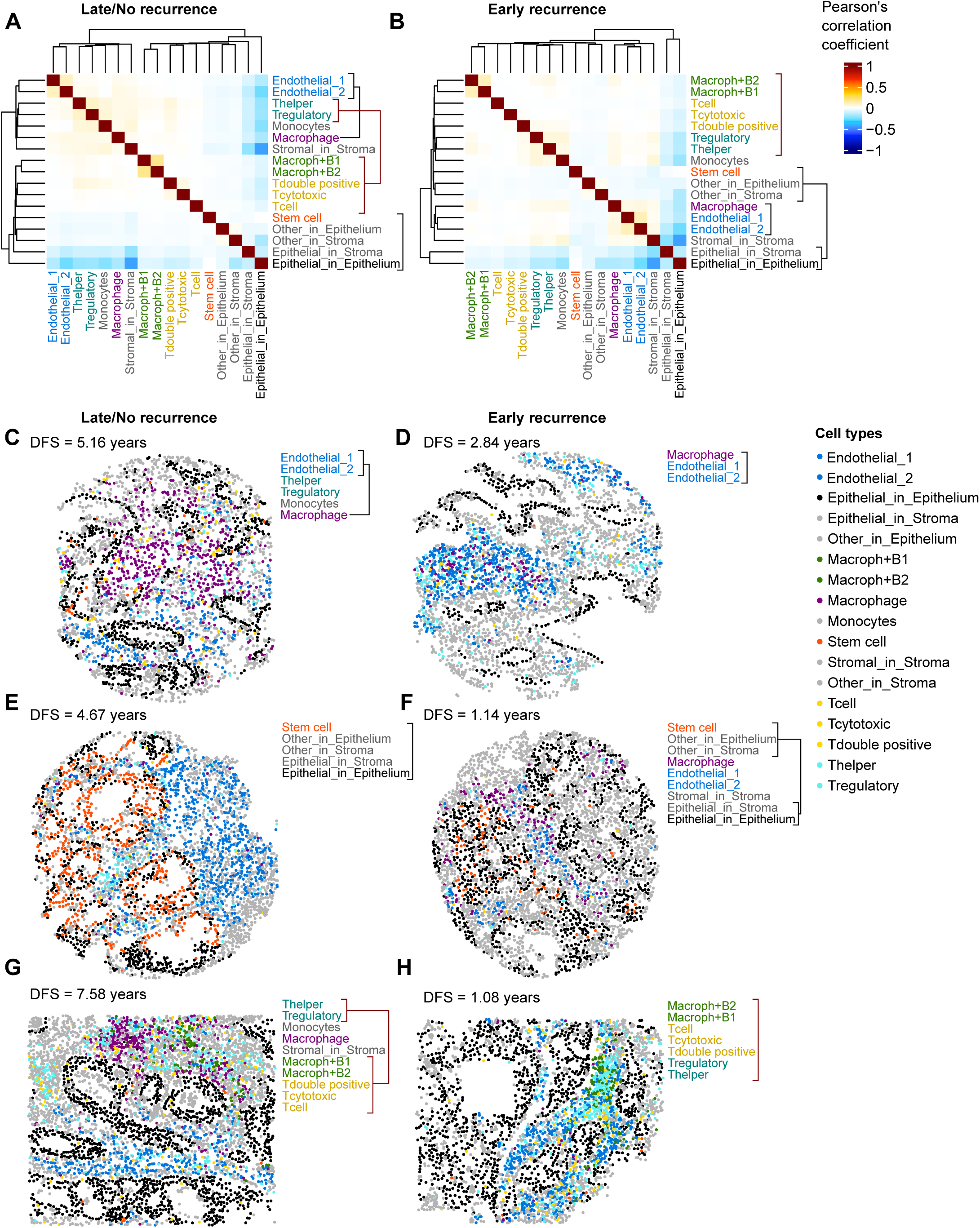
Cell neighbourhood differences between early and late/non-recurring patients. Pairwise spatial correlation of neighbourhood probabilities between different cell types in the group of late/non (A) and early (B) recurrent cores. (C, D) The example of cores from different recurrence groups that are demonstrating the differences in the macrophage proximity to the two types of endothelial cells. (E, F) The example of cores from different recurrence groups that are demonstrating the differences in integrity of epithelium and stem cells within it. (G, H) The example of cores from different recurrence groups that demonstrate the differences in composition of immune hot spots.

A first feature we detected was the proximity of macrophages to endothelial cell groups in the cores of patients with early recurrence (**Figure 3B, D**) compared to the group of cores from late/non-recurring patients (**Figure 3A, C**). This suggests that macrophages spatially associated with blood vessels may promote angiogenesis and be better positioned to induce more frequent intravasation events that lead to early metastatic recurrence ^11^.

The second feature that we found in early recurring cores were stroma-associated stem cells spatially separated from the epithelium (**Figure 3B, F**). The colon epithelium comprises epithelial cells expressing markers CDX2+ and PCK26+ (Epithelial_in_Epithelium cells) tightly clustered together. Typically, stem cells should remain within epithelium layer and not extend into stromal regions. In early recurring samples, stem cells were dispersed away from Epithelial_in_Epithelium cells. In contrast, in late/non-recurring samples, stem cells were spatially closer to Epithelial_in_Epithelium cells (**Figure 3A**), suggesting better preservation of the epithelial crypt structure’s integrity (as shown in **Figure 3E**) compared to early recurrence samples (**Figure 3F**).

The third feature of early recurring samples was a frequent colocalization of Tregulatory and Thelper with B cell-associated macrophages. These were also in proximity to Tcytotoxic, Tdouble positive and Tcell lymphocytes (**Figure 3H**). In contrast, we found that Tregulatory and Thelper cells in late/non-recurring samples were mostly excluded from this close proximity arrangement (**Figure 3G**). The conglomeration of B cells with Tregulatory, Thelper, Tcytotoxic and Tcell lymphocytes suggests the formation of immune ‘hot spots’, some of which possibly resemble tertiary lymphoid structures (TLS) that can play critical roles in antitumor immunity ^12,13^. The identification of ‘hot spots’ also in early recurring patient samples suggests that factors other than antitumoral defence mechanism may play a more prominent role in a subset of early recurring patients.

### Stroma enrichment with high numbers of T regulatory and T helper cells are prominent in tumour tissue of late/non-recurring patients

Next, we analysed the frequency of each cell type within stroma, epithelia and the entire core area across individual patients from all four clinical cohorts (**Table S1**) and how these relate to prognosis (**Figure 4A-C, Figure S2**). Our results showed a substantial, significant increase in the percentage of Tregulatory and Th cells in the late/non-recurring samples in both stroma (**Figure 4A**) and across the entire core (**Figure S2**). This finding additionally validated our previous discovery in a distinct, smaller patient cohort of 5-FU-treated stage III CRC patients where a Cox regression model predicted similar trends ^14^. The percentage of epithelial cells dissociated from the epithelium and located in the stroma was also found to be higher in the early recurring samples (**Figure S2**), suggesting a higher propensity of epithelial cells for EMT is prognostic of early recurrence. We also observed that overall stem cell frequency was not indicative of recurrence (**Figure 4B**).

**Figure 4.**
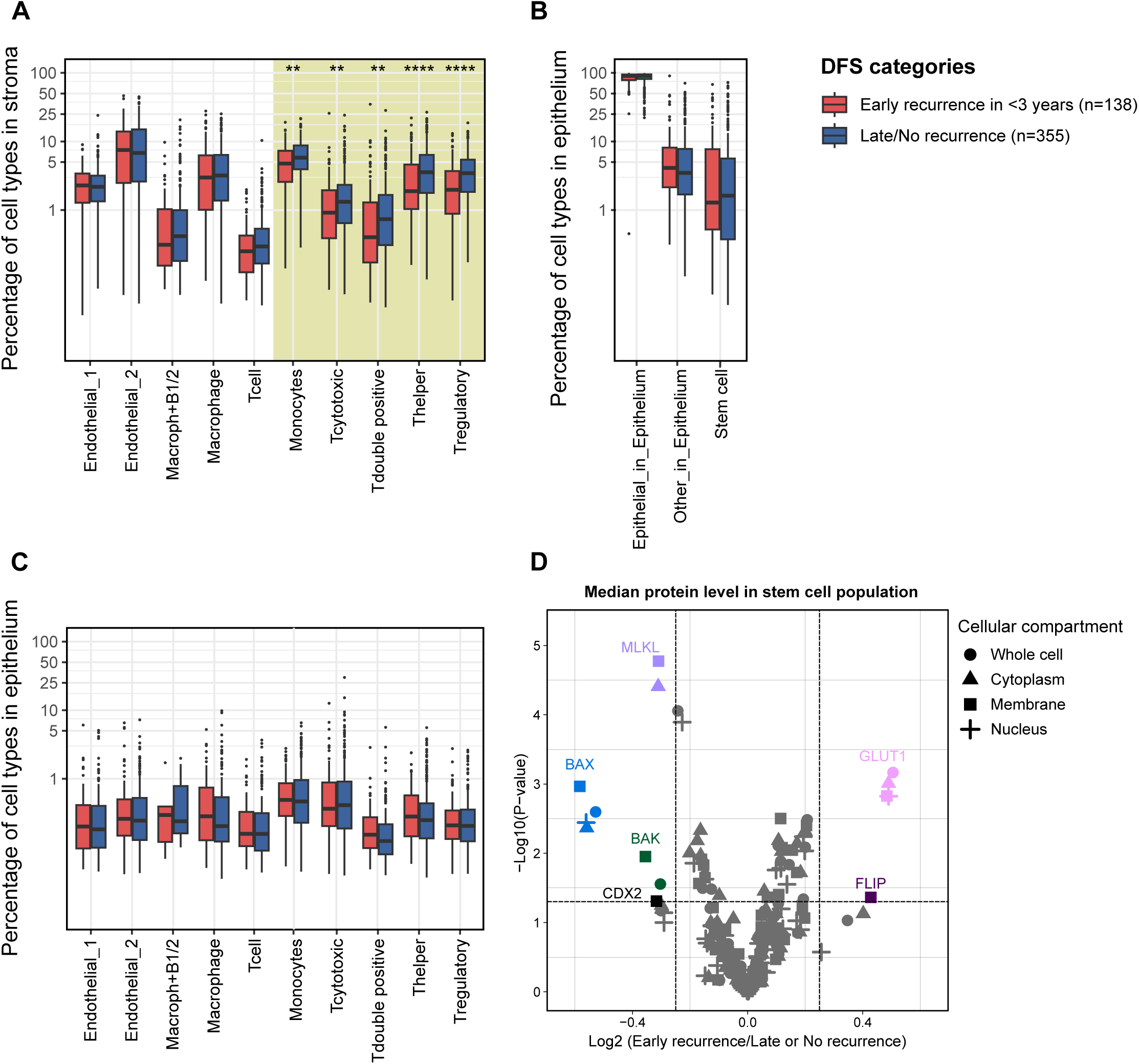
Prognostic cell state markers found in the stem cell population. (A-C) Cell types frequencies are compared between early and late/non-recurrent groups of patients. Cores from individual patients are combined to calculate the percentage of each cell type over stroma and epithelium. Boxplot represents interquartile range (IQR) with the median, 25% (Q1) and 75% (Q3) quantiles as middle, lower and upper hinges, respectively. Lower and upper whiskers are Q1-1.5*IQR and Q3+1.5*IQR, respectively. P-values of the mean cell type frequency compared by the Wilcoxon test are indicated for significant difference between early and late/non-recurrent groups of patients (* P<0.05, ** P<0.005, *** P<0.0005, **** P<0.00005). Differential marker expression analysis in stem cell population (D) performed on the single core bases combining single cell expression into the median signal over the stem cell population of each core individually. Significance P-value cutoff at 0.05 is indicated by horizontal dashed line and log-fold change cutoffs at 0.25 are indicated by a vertical dashed line.

### Stem cells in early vs. late/non-recurring samples show a prominent difference in protein profile

Even though the frequency of CSCs was not prognostic, we hypothesised that the specific biology of CSCs may predict their propensity as propagators of metastasis when resident in the primary tumour. To explore this hypothesis, we compared the expression profiles of CSC between early and late/non-recurring samples using a differential expression analysis of quantified protein markers. The median expression of cell ‘state’ markers of the stem cell population in individual cores were calculated for four cellular compartments (whole cell, cytoplasm, membrane and nucleus). In this analysis, we considered individual tissue cores (2-3 core per patient) rather than an aggregated (i.e. patient) level to account for intratumoral heterogeneity as every core was taken from different sites of the patient’s tumour tissue. This analysis revealed 6 protein markers that were significantly differently expressed and could be considered to constitute a prognostic CSC signature (**Figure 4D**). We detected overexpression of two markers in stem cells of early recurring samples. One of these proteins is a marker of cell metabolism that was previously found to be associated with tumour progression. This protein, glucose transporter 1 (GLUT1), is known to contribute to the Warburg effect in cancer cells ^15^. The second overexpressed protein was cellular-FLICE inhibitory protein (FLIP) that blocks extrinsic apoptosis by interfering with the activation of caspase-8 at death-inducing signalling complexes (DISCs) ^16^ upon death receptor stimulation by death ligand-expressing immune effector cells. Among down regulated proteins, we detected the pro-apoptotic proteins BAK and BAX that are the key components of the intrinsic apoptotic pathway that ensure mitochondrial outer membrane permeabilization (MOMP) ^17–20^, as well as mixed lineage kinase domain-like (MLKL) pseudokinase that controls necroptotic cell death resulting from plasma membrane permeabilization ^21,22^. Finally, we detected the loss of caudal-related homeobox transcription factor 2 (CDX2), a protein that regulates homeostasis of intestinal epithelial cells and cell adhesion. Importantly, such profiles were not detected in epithelial ‘non-stem’ cancer cells where the differential protein profile lacks MLKL, GLUT1 and FLIP signatures (**Figure S3**). This analysis revealed that stem cells found in primary tumour samples from early recurring patients do not only show a different localisation pattern but are also primed to promote EMT and resist apoptotic and necroptotic cell death signalling as well as metabolic stress.

Limited differential signatures were detected in a few other prominent cell types (**Figure S3**). For example, Thelper, Macrophage and Tdouble positive CD4+CD8+ cells showed CASP8 downregulation in early recurring samples. This may indicate resistance to extrinsic apoptosis in these cells types. Th2 cells ^23^ and M2 macrophages ^24^ are known contributors to immunosuppression, and Th2 polarisation can be triggered by Tdouble positive CD4+CD8+ cells ^25^. Thus, cell death resistance in these cell types could be indispensable for tumour propagation. Collectively, however, the stem cell population offered the best potential for development of a prognostic signature.

### A deep neural network delivers a highly prognostic model of recurrence in stage III CRC patients

Deep neural network (DNN) based approaches offer indispensable capabilities and solutions to the challenges in clinical oncology and medical data analysis. We next used DNN to build a prognostic model for early CRC recurrence based on the markers differentially expressed in stem cells in early recurring versus late/non-recurring patients. The successful DNN model was identified in Campion/Challenger approach based on 5 differentially expressed markers (cytoplasmic BAX, MLKL, FLIP, GLUT1 and whole cell CDX2). We trained our DNN with 700 patient cores from 3 discovery cohorts (HV, MSK, Taxonomy) (**Table S4**). During training, the data set was split between training and validation sets at patient level. In each model, variant hyperparameter optimization around the repeated k-fold method was employed to generate multiple balanced splits between training and validation data sets assisting the network’s prediction accuracy by F1-score of early recurrent class of cores in the validation subset (**see Methods**). The champion network was able to capture significant differences in the recurrence dynamics between groups of early and late/non-recurring patient cores in the resulting training (**Figure 5A**) and validation cohorts (**Figure 5B**) that were subsequently verified in the independent validation cohort (RCSI, **Figure 5C**).

**Figure 5.**
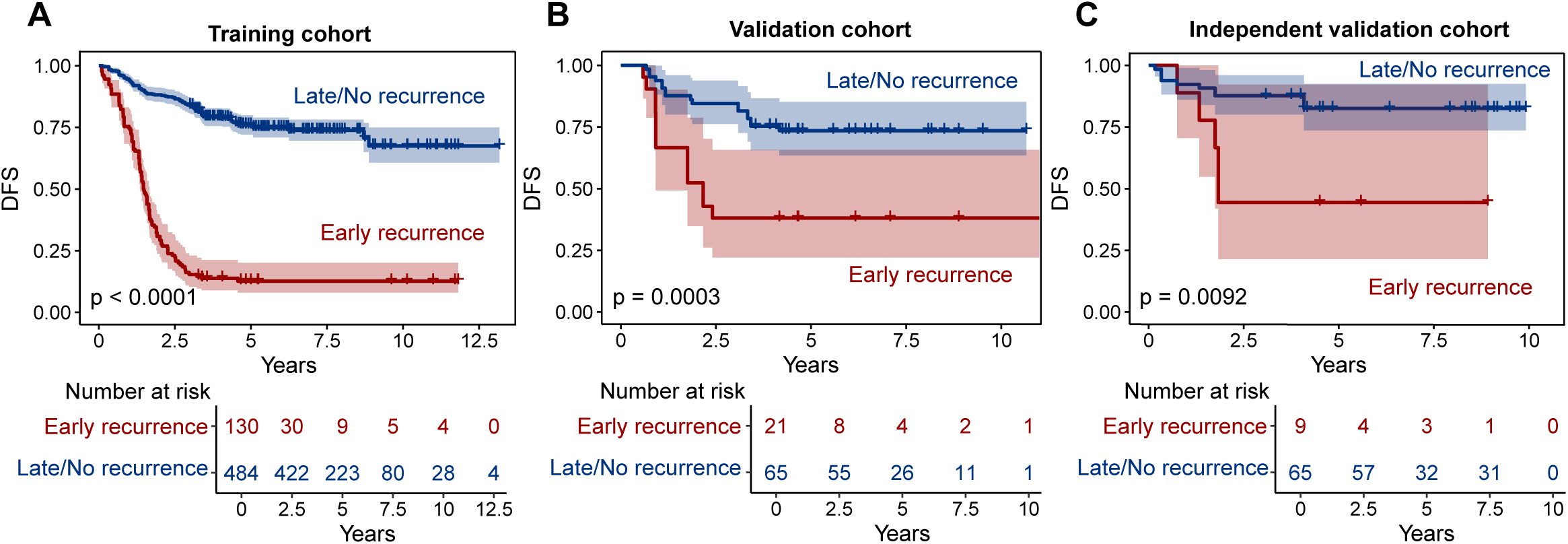
Accurate prognostic model is established by DNN based on 5 protein markers. High accuracy of DNN model based on 5 markers (cytoplasmic BAX, MLKL, FLIP, GLUT1 and whole cell CDX2) in the training (A), validation (B) and independent validation (C) rounds demonstrated in significant difference between early and late/non-recurrent groups of cores on Kaplan-Meier plots.

Finally, clinicopathological data were incorporated to explore whether this discriminated further the recurrence groups. We performed univariate Cox regression analysis identifying clinical parameters in all four cohorts that associated with increased hazard ratio (**Table S5**) and tested them in a multivariate Cox regression model (**Figure 6A**) to account for potential covariates such as age and sex. We identified N staging as the key parameter that further improved prognostic accuracy when combined with predictions of our DNN stem cell model (**Figure 6B**), delivering a new and powerful concise protein signature of disease recurrence.

**Figure 6.**
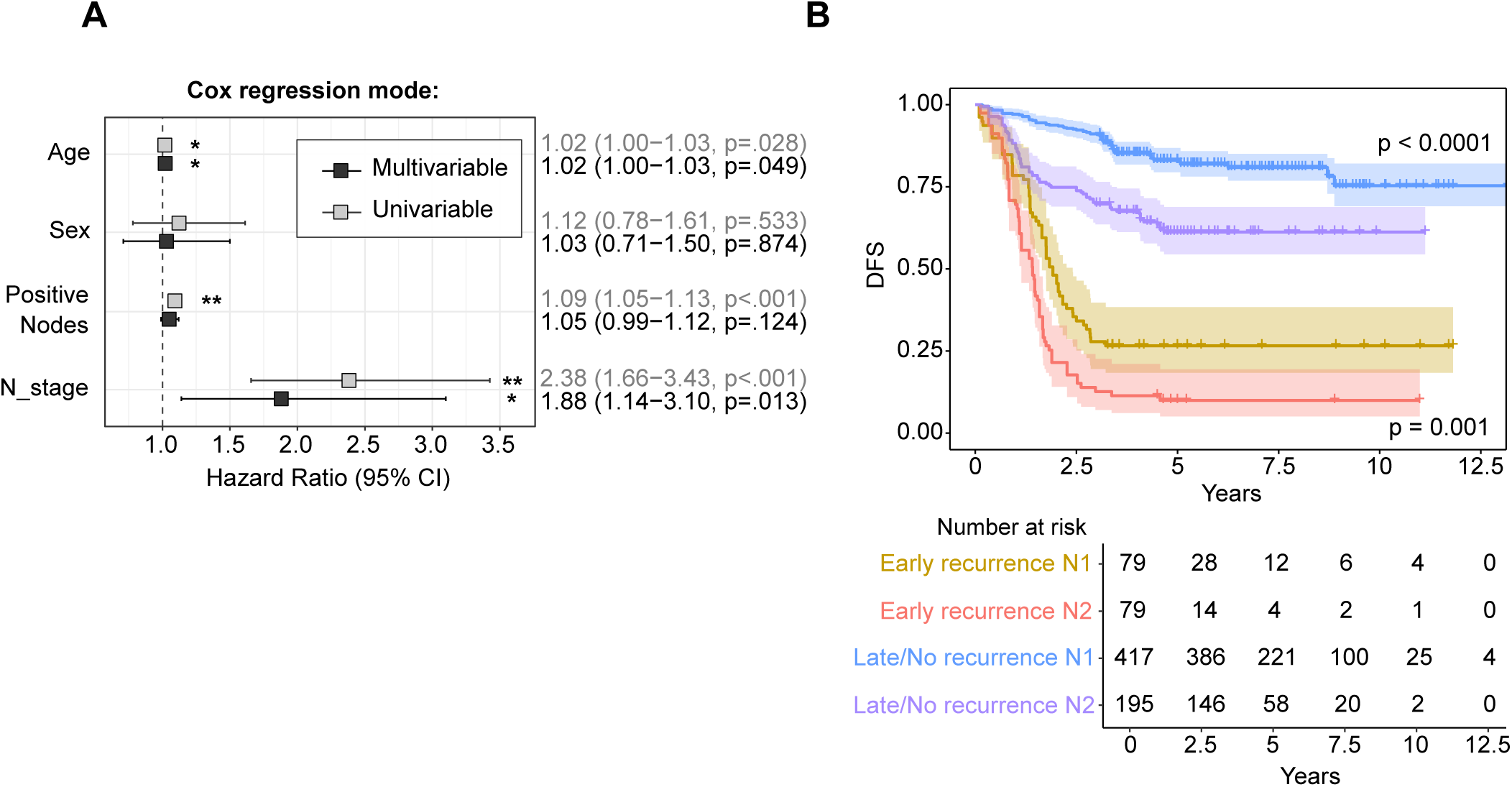
Improved accuracy of prognostic model with inclusion of regional lymph nodes (N) stage. (A) The Cox’s proportional hazards regression analysis of clinical parameters based on multivariate and univariate model. (B) The improved accuracy in the prognosis with combination of DNN model and N stage information in the four cohorts combined.

## Discussion

The results of this study identify a stem cell-specific protein profile that correlates with tumour relapse, highlighting several key proteins that deliver accurate prognostic predictions from the primary tumours of patients diagnosed with stage III CRC and treated post-surgically with adjuvant chemotherapy. In addition, this study highlights several patterns of cell proximity arrangements, including immune, endothelial, and stem cells present in primary tumours that are associated with early tumour recurrence.

We first developed a cell classification pipeline that allowed unbiased identification of 15 cell types over the multiplexed and processed TMA slides. The cell type frequency analysis shows the association of higher stromal abundance of regulatory, helper, cytotoxic and double positive T cells with better prognosis. Next, we analysed cell neighbourhoods to find characteristic spatial proximity arrangements that specifically discriminate early from late/non-recurring patients. By including all cell types found in our neighbourhood analyses, the scope of our search was broad, ranging from identifying spatially organised antitumoral lymphocyte arrangements ^26^ and TLS-like structures ^13,27^ to cellular neighbourhoods that promote macrophage-fibroblast communities ^28,29^ and migration events of CSCs ^30^. As a result, we identified three prominent features: Firstly, we found that early recurring cores more frequently include proximity arrangements of macrophages among groups of endothelial cells. This indicates the association of macrophages with blood vessels that, considering recent findings from other laboratories ^11^, could indicate the initiation of intravasation events that lead to early metastatic recurrence. In previous studies, such perivascular macrophages were found to be a cause of local transient hyperpermeability of tumour vasculature ^31,32^. In turn, this permeability increase may lead to transendothelial migration of cancer cells during intravasation ^33^.

Secondly, we found immune ‘hot spots’ in some early recurring cores, specifically composed of Tregulatory, Thelper surrounded by Macroph+B1, Macroph+B2, Tcytotoxic, Tdouble positive and Tcell. Such ‘hot spots’ can participate in antitumor immunity ^12,13^ and reverse immunosuppression. Indeed, immunosuppression has already been associated with intratumoral immune ‘hot spots’ that include high content of Tregulatory together with B-cells; this occurrence, similar to our findings, was linked to unfavourable outcomes, for example, in lung squamous cell carcinoma (LUSC) ^34^. Important to note that this effect did not correlate with the overall total content of Tregulatory or Thelper across stroma or entire tissue core. In fact, the high percentage of Tregulatory and Thelper cells across whole stroma or core areas was rather associated with late and non-recurring patient samples (**Figure 4A, Figure S2**).

Third, we established that the normal epithelial crypt structure is better preserved in the cores of late/non-recurring patients. Here, stem cells are better integrated within the crypt and do not extend into the stroma of the tissue. This may indicate that stem cells from late/non-recurring patients are less prone to EMT events. Moreover, we observed that the percentage of epithelial cells located in the stroma (Epithelial_in_Stroma, **Figure S2**) was higher in the early recurring samples. This may indicate that metastatic progression may involve both stem and differentiated epithelial cells as cells of origin ^30,35^.

Next, we established a proteomic signature that distinguished stem cells found in early recurring cores from the stem cells in late/non-recurring samples. The most overexpressed proteins in stem cells in early recurring core were GLUT1 glucose transporter and FLIP, a regulator of the extrinsic apoptotic pathway ^16^ and necroptosis ^36^. Among downregulated proteins we found BAK and BAX proteins required for MOMP during intrinsic and BID-dependent extrinsic apoptosis ^17–20^, MLKL pseudokinase, an activator necroptotic cell death ^21,22^, and CDX2 which stimulates cell adhesion in intestinal epithelial cells. Thus, the stem cells from early recurring patients appear to have a unique survival advantage, making them resistant to the most prominent cell death pathways: death receptor-mediated/extrinsic apoptosis, intrinsic apoptosis and necroptosis. Through upregulation of GLUT1, these cells may also increase glucose influx rewiring their metabolism to promote growth and proliferation ^15^ and trigger cell migration and malignant transformation through loss of CDX2 which has previously been identified to be associated with poor prognosis ^37^.

Finally, we trained a DNN model to recognize this stem cell signature based on five differentially expressed proteins: BAX, MLKL, FLIP, GLUT1 and CDX2. Our model showed very high prognosis performance separating survival trends for the group of early and late/non-recurring samples, with an exceptionally high significance level, p<0.0001. Furthermore, the DNN model could be further enhanced by inclusion of N staging as clinicopathological parameter, delivering high hazard ratios in our patient cohort.

The ultimate goal for further clinical implementation of our prognostic signature is the establishment and calibration of the robust pipeline (**Figure S6**) compatible with the multiplex technologies available for clinical use, for example, such as ZEISS Axioscan 7 and Leica Aperio VERSA for imaging and quantification of 5 markers of the prediction signature as well as segmentation marker such as PCK26 and stem cell marker ALDH1 or other alternative stem markers such as leucine-rich repeat-containing G-protein-coupled receptor 5 (LGR5), CD44 or Musashi1 (MSI1). Additionally, the inclusion of the simultaneous brightfield image, as on ZEISS Axioscan 7 and Leica Aperio VERSA platforms, can illuminate the necessity of fluorescent staining for segmentation marker as the Hematoxylin and eosin staining (H&E) can provide accurate segmentation of the cytoplasm regions as well as epithelium with the appropriate image processing pipeline. Additionally, the use of larger tissue samples, currently relevant to the clinical setting, can further improve the accuracy of the prediction method.

Overall, our results propose a clinically promising prognostic tool based on a five-protein stem cell signature. Compared to currently proposed transcriptomic based signatures ^38^ our prognostic signature exhibits superior discriminative ability which is demonstrated by a substantial separation between risk groups. Moreover, our five-protein signature markers not only predict stem cell chemotherapy resistance and therefore tumour recurrence but also suggest potential therapeutic strategies. For instance, this approach could guide combinatorial treatments at high risk of chemoresistance, such as incorporating small molecule inhibitors targeting FLIP (currently in discovery phase) and GLUT1 (already under preclinical trial evaluation).

## Methods

### Clinical cohorts description

This study was performed in accordance with ethical guidelines for clinical research with the approval of the ethics committee for each institution. Formalin-fixed, paraffin-embedded primary tumour tissue sections were obtained from four different cohorts. For “RCSI” cohort, tissues were provided by the Beaumont Hospital (RCSI, Ireland) Colorectal Cancer Biobank with written consent provided by all patients. Institutional ethical approval was granted by the Beaumont Hospital Research and Ethics Committee (References 08/62 and 19/46). For “Taxonomy” cohort, tissues were supplied by the Queen’s University Belfast (QUB, UK) Department of Pathology with written consent provided by all patients and institutional ethical approval granted (NIB12-0034). For “HV” cohort, samples were collected at Huntsville Clearview Cancer Center (CCI, US) under granted institutional ethical approval (IRB/CC1 Colon 01) and, for “MSK” cohort, tissue were provided by Memorial Sloan Kettering Cancer Center (MSK, US) under ethical agreement (IRB/WA0497-09). All patients included in the study underwent surgery for histologically proven stage III CRC cancer. Only patients with adjuvant and without neoadjuvant therapy were included, and only cases for which we had access to tissue blocks on the original resection and clinical follow-up information were included. Patients’ exclusion criteria were as follows: postoperative mortality within 30 days; a limited follow-up period of less than 3 years in cases without recurrence; synchronous multiple cancers; positive surgical margins; concomitant inflammatory bowel disease; and familial adenomatous polyposis or previous malignancy within 5 years. A clinical summary of all cohorts can be found in **Table S1**.

### The Patient and Public Involvement statement

Because this was a retrospective study of archived samples and data, patients or the public were not involved in the design, or conduct, or reporting or dissemination plans of our research.

### Differential expression analysis

The median expression of cell ‘state’ markers of the stem cell population in individual cores were calculated for four cellular compartments (whole cell, cytoplasm, membrane and nucleus).

In this analysis we considered individual cores rather than patients to account for intratumoral heterogeneity considering that every core was taken from different sites of patients’ tumour tissue. Despite our normalization pipeline ^39^ created to equalize the batch effects for every protein across several TMA slides, some proteins may remain affected. Therefore, to find robust protein markers which are independent of batch effects, and that differentiate early from late/non-recurring stem cell populations, the differential expression analysis was conducted with and without consideration of TMA slide related batch effects. Then the resulting strong significant difference in expression detected in the analysis without corrections and confirmed in the analysis with corrections were then considered for prognostic markers selection (**Figure 4D**). Thus, the initially normalised intensities for these selected markers can be confidently used further in the prediction model training that accounts for small variance that may be present due to the batch effects.

### Deep neural network (DNN) training

We trained our DNN with 700 patient cores from three cohorts (HV, MSK, Taxonomy cohorts) (**Table S4**). During training the data set was split between training and validation sets at patient level. We used repeated 8-fold cross-validation method (20 repeats) to assess generalisation ability across multiple data splits. In each split the general proportional balance of early and late/non-recurring patients we preserved for training and validation sets. Each training event was started with initial learning rate that gradually decay over training steps progression (epoch) with the given decay rate. To avoid internal covariate shifts in the DNN layers the layer weights batch normalisation was applied. A class weight hyperparameter was applied to compensate for imbalance class representation (early and late/non-recurring cores) and was also subject for optimisation. Feature importance was assisted with L1-norm value in the first layer of the DNN. The validation performance was evaluated by calculating the F1-score for predicting early recurrence, accuracy, precision, specificity and recall. We optimised the DNN hyperparameters to maximise the F1-score in validation set using Champion/Challenger approach. The hyperparameters considered in Adam optimisation ^40^ procedure were: Initial learning rate, learning decay rate, class weight, activation function, number of hidden layers, size of each hidden layer, dropout rate, penalties for L1 and L2 regularisation. We implemented the early stopping patience principal to prevent over-training, which terminated the training when the validation F1-score was no longer improving for 500 consecutive epochs. The early stopping was allowed only in the stable learning phase that begins at 20th epoch. Finally, the champion DNN was validated in the independent validation cohort (RCSI cohort) (**Table S4**). The neural network was implemented in Julia v1.9.3 ^41^, using Flux.jl v0.14.6 ^42^ for architecture design and training, whilst MLUtils v0.4.3 handled data preprocessing operations including observation shuffling and train-validation splitting whilst MKL v0.6.1 accelerates matrix operations through Intel’s Math Kernel Library backend.

### Statistics

Disease-free survival (DFS) was defined as the time from surgery to the date of local recurrence or relapse of CRC, distant metastasis, or death from an underlying disease. Survival characteristics were described using the Kaplan-Meier method and compared using the log-rank test. Hazard ratios were calculated using the Cox proportional hazard model. All conventional statistical analyses were performed in the R environment ^43^. The level of statistical significance was set at *P* < .05. Wilcoxon tests were performed using ggpubr v0.3.0 package, differential expression analysis was done with the moderated t-test with Benjamini-Hochberg FDR adjustment in the *limma* package (v3.40.6) ^44^ and plots visualisation were performed with the ggplot2 v3.3.0 package ^45^. The co-localization neighbourhood probabilities between different cell types were calculated with the hoodscanR package in R ^8^. Hierarchical clustering was implemented in Julia v1.9.3 ^41^, using Clustering v0.15.8 where Ward’s method with the Euclidean linkage was implode. MKL v0.6.1 were used for acceleration of matrix operations through Intel’s Math Kernel Library backend. LDA runs were performed with R package MASS v7.3-51.5 ^46^.

## Acknowledgement

We thank Dr Andreas Lindner for support with cell classification pipelines.

## Authors contribution

AS, SaCh, MS, MSt, BK, EMcD, EPO’C, JFG, SSMD and MA were involved in methodology, data validation, and curation. SaCh, EMcD, JFG, and FG performed the Cell DIVE processing as well as cell segmentation and single-cell quantification. AS performed cell type classification, statistically analysis and study investigation. JPB, NMcC, DAmcN, JS, CF, NU, JHMP, and DBL were involved in clinical sample acquisition and data collection. EMcD conducted the sample imaging and image processing. AS and JHMP wrote the manuscript. DBL, FG, and JHMP were involved in funding acquisition. JHMP supervised the study. All authors edited and revised the manuscript text.

---

***Table S1.*** *Baseline table with characteristics of HV, MSK, RCSI and Taxonomy clinical cohorts.*

***Figure S2. Prognostic cell state markers found in the stem cell population.*** *Cell types frequencies are compared between early and late/non-recurrent groups of patients. Cores from individual patients are combined to calculate the percentage of each cell type over tissue core area. Boxplot represents interquartile range (IQR) with the median, 25% (Q1) and 75% (Q3) quantiles as middle, lower and upper hinges, respectively. Lower and upper whiskers are Q1-1.5*IQR and Q3+1.5*IQR, respectively. P-values of the mean cell type frequency compared by the Wilcoxon test are indicated for significant difference between early and late/non-recurrent groups of patients (* P<0.05, ** P<0.005, *** P<0.0005, **** P<0.00005).*

***Figure S3. Differential markers found in other cell populations.*** *The volcano plots show the results of differential marker expression analysis in all classified cell populations performed on a single-core basis, combining single-cell expression into the median signal over the given cell type population in each tissue core individually. Significance P-value cutoff at 0.05 is indicated by a horizontal dashed line and log-fold change cutoffs at 0.25 are indicated by vertical dashed lines.*

***Table S4.*** *Baseline table with clinical characteristics for patients with identifiable stem cell signature from HV, MSK, RCSI and Taxonomy clinical cohorts used in the training and validation of the DNN model.*

***Table S5.*** *Univariate Cox proportional hazard analysis.*

***Figure S6***. ***The overview of the plausible adaptation concept of the acquisition and application of the prognostic signature for the prediction of CRC recurrence in stage III CRC samples.** The pipeline begins with appropriate image segmentation to identify the epithelium and stem cell cytoplasm regions at the single cell level. As an example, we used PCK26 staining alone for epithelium and single cell detection, however, it can be complimented or substituted by H&E staining based segmentation pipeline. Next, stem cells are identified through the ALDH1 fluorescent signal, resulting in the Stem_mask generation. Finally, the overlay of single cell epithelial Cytoplasm_mask with Stem_mask delivers the single cell Stem_Cytoplasm_mask. This mask is used to acquire the mean pixel intensities of the fluorescent signal for 5 proteomic markers of the prognostic signature over the area of individual stem cells. These quantities are further agglomerated in the median protein signal across all stem cell populations for the individual core sample, thus creating the final core level 5 protein prediction signature. This signature input into our DNN prediction network that delivers the prognosis prediction.*

