## Supplementary Methods for "A neural network model delivers a highly prognostic protein signature in cancer stem cells that identifies relapse in stage III colorectal cancer patients"

**Cell type classification pipeline**

We developed a pipeline for cell type identification for high-plex single cell data analysis in stage III tumour tissue. The boundaries between stroma and epithelium of the tissue cores were established by creating binary mask based on PCK26 expression (1). The cell analysis pipeline begins with the recognition of epithelial CDX2^+^, PCK26^+^ and AE1^+^ positive cells (‘cancer cells’) using single-cell markers fluorescent signal intensity distributions approximated by kernel density estimation, followed by automated detection of the extremes within the distribution span (**Figure 2A**). The minimum of probability distribution between two local maximums identified a binary classification threshold on an individual TMA slide basis.

The same threshold detection algorithm was used to classify immune cells. A subset of the CDX2^-^/PCK26^-^/AE1^-^ cell population was considered, firstly, for CD3 based threshold estimation to determine the population of CD3^+^ immune cells, and secondly, to categorise CD8^+^ and CD4^+^ immune cells among the CD3^+^ immune cell population. The distribution of the FOXP3 signal across the CD3^+^CD4^+^ cell population to characterise regulatory T cells (Tregulatory) was less discrete and therefore required a different approach to approximate thresholds. Therefore, we used a Gaussian mixed model to fit the FOXP3 signal density distribution. The position of the distribution’s mean characteristic for FOXP3^-^ cell population was estimated from the Gaussian model fit on the subset of CD3^+^CD4^-^ immune cells that are generally known for the lack of FOXP3 expression. As such, the final threshold was recognised as the intersection of the two Gaussian models for FOXP3^-^ and FOXP3^+^ cell populations in the original CD3^+^CD4^+^ cells subset. Collectively, this pipeline identified the population of epithelial cancer cells, Monocytes (CD3^-^CD8^-^CD4^+^), T regulatory (CD3^+^CD4^+^FOXP3^+^), T helper (CD3^+^CD4^+^FOXP3^-^), double negative T cells (Tcell CD3^+^CD8^-^CD4^-^FOXP3^-^), cytotoxic T cells (CD3^+^CD8^+^) and double positive T cells (CD3^+^CD4^+^CD8^+^) (**Figure 2A**).

The classification of other cell types such as endothelial, macrophage, and B cells required the combination of several markers that may exhibit non-convex distributions. We therefore employed divisive hierarchical clustering as a method that provides unparalleled accuracy but required increased computational resources. This method is suitable for processing of a smaller subset of randomly selected, unclassified cells from the remaining cells in the stroma of the TMA slides. We used cell marker profiles to build a classifier by means of a linear discriminant analysis (LDA) approach (2). The LDA approach was applied to finalise the classification over remaining unclassified cell subsets in the TMA slide. Thus, we built classification models for macrophages (Macrophage), B cells (Macroph+B1/B2) and endothelial cells (Endothelial_1/2) by first performing hierarchical clustering with 7 markers (CD20, CD31, CD34, CD4, CD68, COLIV and SMA) on a subpopulation of 90000 randomly selected CD3^-^/CD8^-^/AE^-^/PCK26^-^/CDX2^-^ unclassified cells. The optimal clusters composition was estimated based on the average silhouette method (3) for every clustering run individually for each TMA slide. The cluster with high CD4 and CD68 expression (CD4↑CD68↑) was assigned the Macrophage cell type. A cluster with a CD31↑CD34↑COLIV↑SMA↑ profile represented ‘Endothelial_1’ cell. A cluster with a CD31↑CD34↓COLIV↑SMA↑ profile represented a second type of endothelial cells, which we termed in our classification ‘Endothelial_2’. Finally, in every TMA slide we detected two signatures of B-cells characterised by high or moderate CD20 expression (CD20↑ and modCD20, respectively), however these signatures were always accompanied by substantial CD68 and CD4 expression. Therefore, two new cell types with CD4↑CD68↑CD20↑ and CD4↑CD68↑modCD20 signatures were identified and named as B cell-associated macrophage populations, Macroph+B1 and Macroph+B2.

Stem cells were identified following the same principles, with the clustering on the fraction of the remaining unclassified cells in the epithelium. Stem cells were recognized by ALDH1↑KI67↓AE1↓ signature following discovery in the remaining subset by LDA model (**Figure 2B**).
