## Supplementary Table S1 for "A neural network model delivers a highly prognostic protein signature in cancer stem cells that identifies relapse in stage III colorectal cancer patients"

| Characteristic | HV<br>N = 161[1] | MSK<br>N = 252[1] | RCSI<br>N = 42[1] | Taxonomy<br>N = 38[1] | p-value[2] |
|---|---|---|---|---|---|
| age | 63 (54, 71) | 59 (50, 70) | 66 (57, 73) | 67 (58, 71) | 0.006 |
| sex | | | | | 0.2 |
| female | 70 (43%) | 133 (53%) | 22 (52%) | 16 (42%) | |
| male | 91 (57%) | 119 (47%) | 20 (48%) | 22 (58%) | |
| N_stage | | | | | 0.3 |
| N1 | 98 (61%) | 163 (65%) | 27 (64%) | 29 (76%) | |
| N2 | 61 (38%) | 89 (35%) | 15 (36%) | 9 (24%) | |
| N3 | 2 (1.2%) | 0 (0%) | 0 (0%) | 0 (0%) | |
| T_stage | | | | | |
| T1 | 4 (2.5%) | 6 (2.4%) | 1 (2.4%) | 1 (2.6%) | |
| T2 | 20 (13%) | 26 (10%) | 2 (4.8%) | 3 (7.9%) | |
| T3 | 124 (78%) | 171 (68%) | 27 (64%) | 27 (71%) | |
| T4 | 11 (6.9%) | 49 (19%) | 12 (29%) | 7 (18%) | |
| Unknown | 2 | 0 | 0 | 0 | |
| Recurrence | | | | | 0.003 |
| Early | 54 (34%) | 57 (23%) | 9 (21%) | 18 (47%) | |
| Late/No | 107 (66%) | 195 (77%) | 33 (79%) | 20 (53%) | |

[1] Median (Q1, Q3); n (%)

[2] Kruskal-Wallis rank sum test; Pearson’s Chi-squared test; Fisher’s exact test; NA
