## Supplementary figures and images for "A neural network model delivers a highly prognostic protein signature in cancer stem cells that identifies relapse in stage III colorectal cancer patients"

### Supplementary Fig. S2

**Supplementary figure**

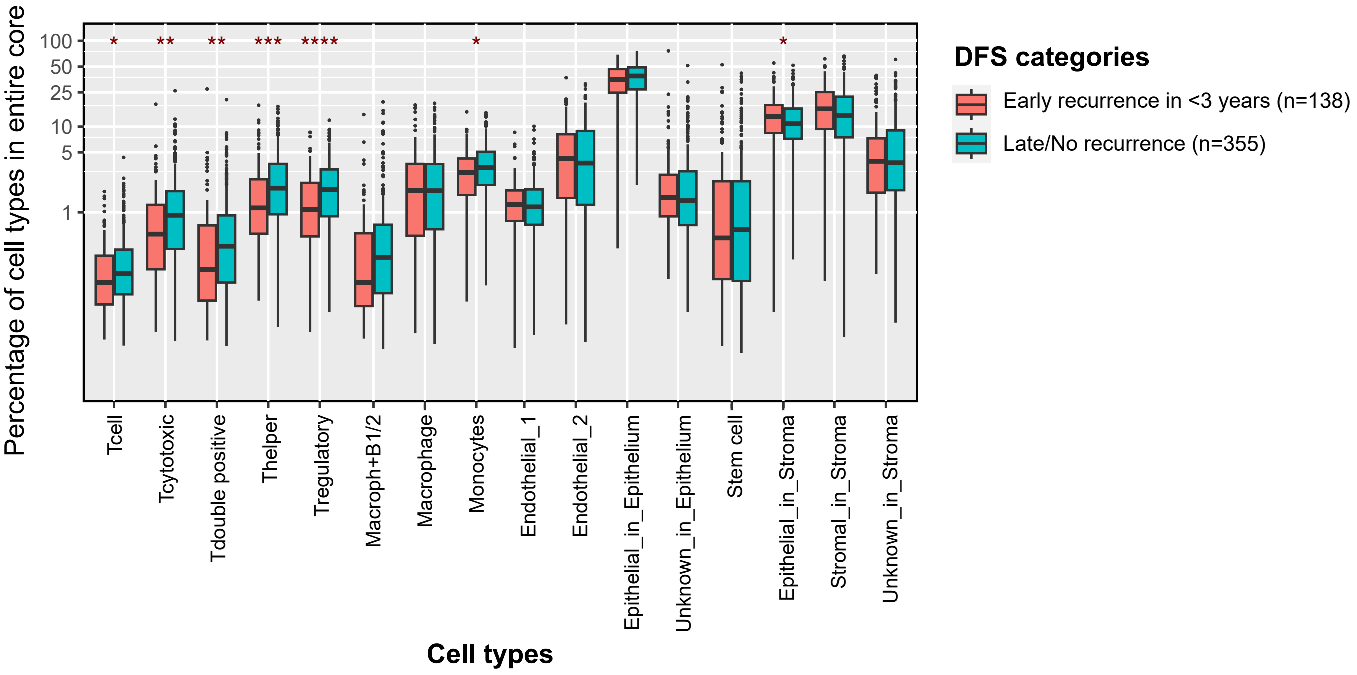

### Supplementary Fig. S3.

**Supplementary figure: Median expression in all cell type populations in each core**

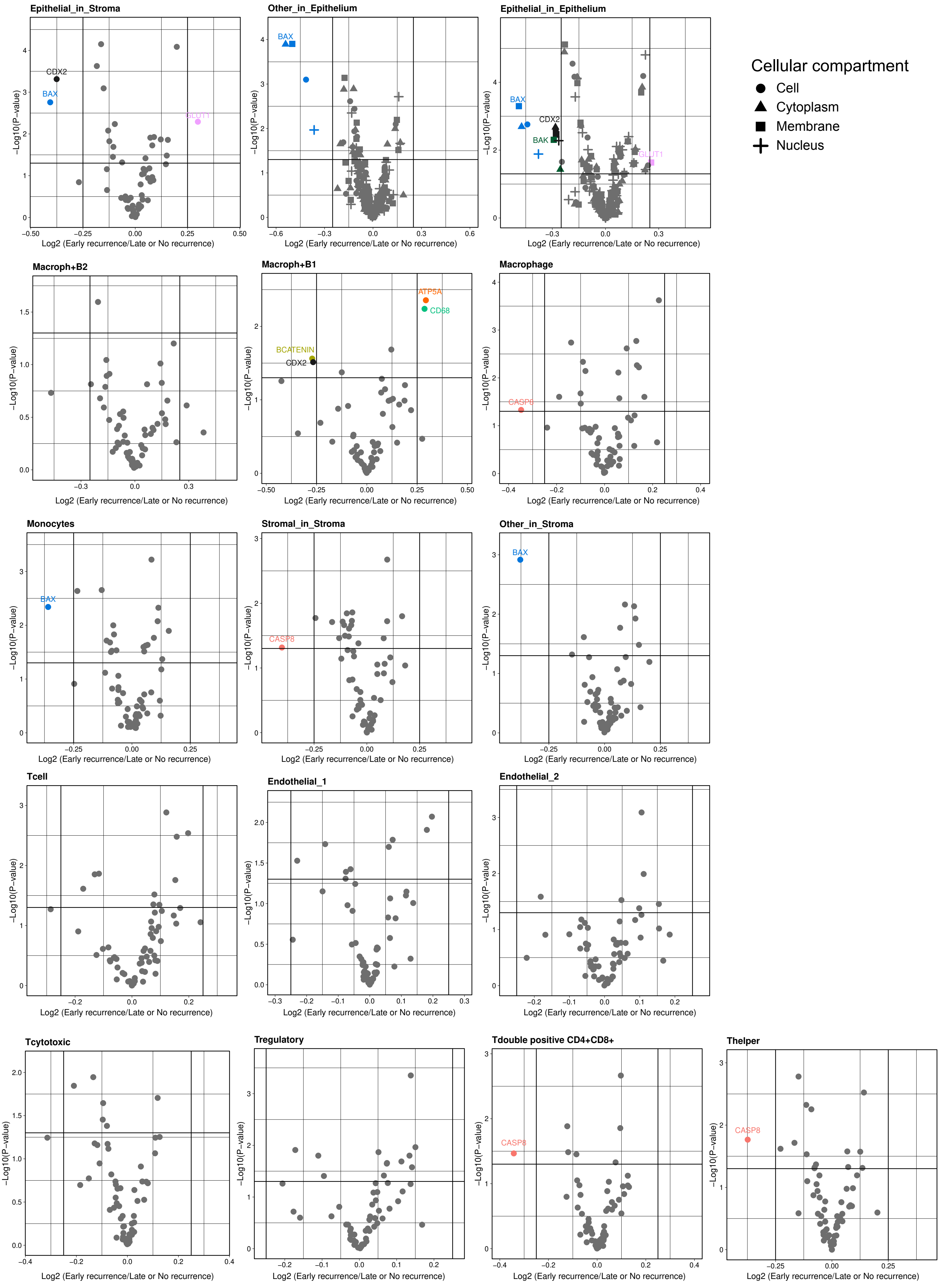

### Supplementary Fig. S6

**Supplementary figure**

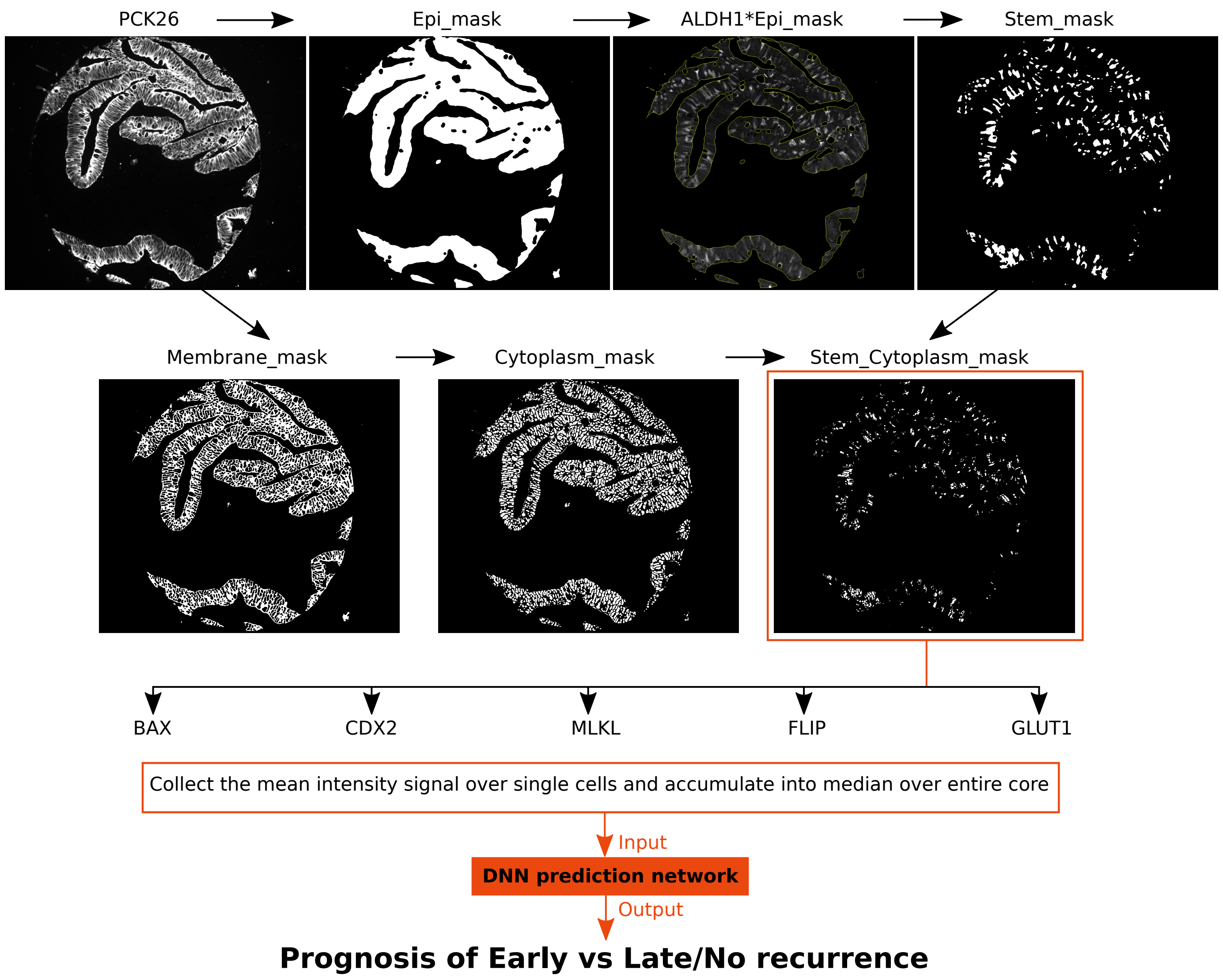
