## Supplementary Table S4 for "A neural network model delivers a highly prognostic protein signature in cancer stem cells that identifies relapse in stage III colorectal cancer patients"

| **Characteristic** | **HV** N = 153[1] | **MSK** N = 183[1] | **RCSI** N = 33[1] | **Taxonomy** N = 27[1] | **p-value**[2] |
|---|---|---|---|---|---|
| age | 63 (54, 71) | 59 (50, 71) | 64 (56, 70) | 67 (57, 70) | 0.069 |
| sex | | | | | 0.6 |
| female | 68 (44%) | 95 (52%) | 17 (52%) | 14 (52%) | |
| male | 85 (56%) | 88 (48%) | 16 (48%) | 13 (48%) | |
| N_stage | | | | | 0.8 |
| N1 | 97 (63%) | 122 (67%) | 22 (67%) | 19 (70%) | |
| N2 | 54 (35%) | 61 (33%) | 11 (33%) | 8 (30%) | |
| N3 | 2 (1.3%) | 0 (0%) | 0 (0%) | 0 (0%) | |
| T_stage | | | | | |
| T1 | 4 (2.6%) | 6 (3.3%) | 1 (3.0%) | 1 (3.7%) | |
| T2 | 20 (13%) | 22 (12%) | 1 (3.0%) | 1 (3.7%) | |
| T3 | 117 (77%) | 126 (69%) | 25 (76%) | 18 (67%) | |
| T4 | 10 (6.6%) | 29 (16%) | 6 (18%) | 7 (26%) | |
| Unknown | 2 | 0 | 0 | 0 | |
| Recurrence | | | | | 0.002 |
| Early | 54 (35%) | 39 (21%) | 6 (18%) | 13 (48%) | |
| Late/No | 99 (65%) | 144 (79%) | 27 (82%) | 14 (52%) | |

[1] Median (Q1, Q3); n (%)

[2] Kruskal-Wallis rank sum test; Pearson’s Chi-squared test; Fisher’s exact test; NA
