## Supplementary Table S5 for "A neural network model delivers a highly prognostic protein signature in cancer stem cells that identifies relapse in stage III colorectal cancer patients"

**Supplementary table:** Univariate Cox proportional hazard analysis

| | beta | HR (95% CI for HR) | wald.test | p.value |
|---|---|---|---|---|
| N_stage | 0.87 | 2.4 (1.7-3.4) | 22 | 2.90E-06 |
| positive_nodes | 0.089 | 1.1 (1.1-1.1) | 21 | 4.40E-06 |
| nodal_count | -0.023 | 0.98 (0.96-0.99) | 6.3 | 0.012 |
| age | 0.017 | 1 (1-1) | 4.8 | 0.028 |
| cea_post_surgery | 0.0038 | 1 (1-1) | 4.8 | 0.028 |
| adjuvant_chemotherapy_cycles | -0.59 | 0.56 (0.3-1) | 3.3 | 0.07 |
| braf_status | 0.46 | 1.6 (0.89-2.8) | 2.4 | 0.12 |
| has_perineural_invasion | 0.4 | 1.5 (0.86-2.6) | 2 | 0.15 |
| has_lymphovascular_invasion | 0.21 | 1.2 (0.9-1.7) | 1.7 | 0.2 |
| tumour_site_1 | 0.14 | 1.2 (0.91-1.5) | 1.4 | 0.24 |
| cancer_type | 0.21 | 1.2 (0.83-1.8) | 1.1 | 0.3 |
| has_extramural_venous_invasion | 0.67 | 1.9 (0.5-7.5) | 0.93 | 0.34 |
| RX | 0.11 | 1.1 (0.88-1.4) | 0.81 | 0.37 |
| p53_status | 0.29 | 1.3 (0.64-2.8) | 0.59 | 0.44 |
| kras_status | -0.38 | 0.69 (0.23-2) | 0.46 | 0.5 |
| sex | 0.12 | 1.1 (0.78-1.6) | 0.39 | 0.53 |
| adjuvant_chemotherapy_regimen | 0.056 | 1.1 (0.88-1.3) | 0.37 | 0.54 |
| Rxtyp | 0.015 | 1 (0.96-1.1) | 0.28 | 0.59 |
| has_mucinous_features | 0.2 | 1.2 (0.59-2.6) | 0.3 | 0.59 |
| MORPHO | 5.90E-05 | 1 (1-1) | 0.23 | 0.63 |
| tumour_size | 0.056 | 1.1 (0.84-1.3) | 0.24 | 0.63 |
| microsatellite_status | -0.19 | 0.83 (0.3-2.3) | 0.13 | 0.72 |
| surgery_year | -0.024 | 0.98 (0.84-1.1) | 0.09 | 0.76 |
| is_micropapillary | 0.31 | 1.4 (0.19-9.9) | 0.1 | 0.76 |
| differentiation | -0.06 | 0.94 (0.52-1.7) | 0.04 | 0.84 |
| histologic_type | -0.071 | 0.93 (0.41-2.1) | 0.03 | 0.87 |
| GRADE | -0.031 | 0.97 (0.55-1.7) | 0.01 | 0.91 |
| race | 0.0083 | 1 (0.84-1.2) | 0.01 | 0.93 |
| tumour_site_raw | 0.0035 | 1 (0.93-1.1) | 0.01 | 0.93 |
| cea_peri_diagnosis | -0.001 | 1 (0.96-1) | 0 | 0.96 |
| tumour_site_2 | -0.0023 | 1 (0.69-1.4) | 0 | 0.99 |
| has_neoadjuvant_chemoradiotherapy | -19 | 4e-09 (0-Inf) | 0 | 1 |
| is_neuroendocrine | -15 | 3e-07 (0-Inf) | 0 | 1 |
| is_signet_ring | -15 | 3e-07 (0-Inf) | 0 | 1 |
| has_medullary_changes | -16 | 1.1e-07 (0-Inf) | 0 | 1 |
| cms_subtype | -22 | 3.5e-10 (0-Inf) | 0 | 1 |
| cris_subtype | -21 | 6.9e-10 (0-Inf) | 0 | 1 |
